# Benchmarking BEAGLE to find optimal parameters for BEAST X

**DOI:** 10.64898/2026.03.10.710534

**Authors:** Samuel Fosse, Sebastian Duchene, Camila Duitama González

## Abstract

Bayesian phylogenetic analyses are notoriously time-consuming, largely because exploring the posterior distribution requires computing Felsenstein’s likelihood. The BEAGLE library is a high-performance computational tool that dramatically accelerates the calculation of such likelihoods by leveraging parallel processing on GPUs, multicore CPUs, and SSE vectorisation. Here we present results from benchmarking a widely popular phylogenetics package, BEAST X, using BEAGLE integration, focusing on how hardware allocation affects running times. We demonstrate substantial differences among BEAGLE settings on real Dengue Virus (DENV) data, both with and without partitioning. Using simulated sequences, we establish guidelines for GPU usage in BEAST X runs. These guidelines can be used for effective resource allocation for empirical analyses and simulation studies.

## I. Introduction

### A. Introduction

Molecular phylogenetic methods use genetic sequence data to make inferences about evolution. Such inferences are largely underpinned by phylogenetic trees. The likelihood of such trees, can be expressed as *P* (𝒟 | 𝒯, 𝒬), where 𝒟 is an alignment of genetic sequence data (i.e. nucleotides or amino acids), 𝒯 is the phylogenetic tree, and 𝒬 is a substitution matrix and its parameters following a Markov chain. Felsenstein’s pruning algorithm is typically used to calculate this likelihood [1].

The field of phylodynamics involves modelling several processes in combination with phylogenetic trees, including epidemiological dynamics and geographic movement (reviewed in [2]). Bayesian methods are particularly popular in phylodynamics because integrating a range of models and prior knowledge is possible through the posterior distribution. For example the posterior distribution *P* (𝒯, *𝒬, µ, θ*|*D*) ∝ *P* (𝒟 | 𝒯, *𝒬, µ*)*P* (𝒯 | *θ*)*P* (𝒬)*P* (*µ*)*P* (*θ*) (where *µ* is the molecular clock rate, *θ* is a demographic parameter, and *P* (𝒟 | 𝒯, *𝒬*) is the likelihood function of a time-tree), can be sampled using popular packages, such as BEAST X [3].

Sampling the target distribution, i.e. the posterior, is not possible with analytical methods. For this reason, Markov Chain Monte Carlo (MCMC) is used to approximate this distribution [4]. Bayesian Evolutionary Analysis Sampling (BEAST) X is a popular program to infer phylogenies using MCMC, and offers a wide array of models for phylogenetic, phylodynamic and phylogeographic inference [3]. BEAST can be computationally intensive, and much has been done to optimise it for the latest hardware architectures [3]. The computation of phylogenetic likelihoods is often the most expensive part of the inference. Since nucleotide sites are assumed to be independent and identically distributed, multi-threading and GPUs can be used to accelerate likelihood computations.

The Broad-platform Evolutionary Analysis General Likelihood Evaluator (BEAGLE) library [5] is an API that provides multi-threaded and GPU-based approaches for evaluating phylogenetic likelihoods. BEAST X implements BEAGLE for likelihood computations, and offers several options to parametrise BEAGLE to optimise running time in function of the available hardware. The speedup factor of parallelising and using GPUs with BEAGLE increases with the number of site patterns, i.e., the number of unique columns in the sequence alignment [5].

Several BEAGLE settings can impact the running time of BEAST. A previous benchmark of BEAGLE [6] evaluated how performance evolves as the number of threads is varied, showing that excessive multi-threading can increase running time. However, this was done in a previous version of BEAST (1.6.1), which predates the current one by more than 10 years. Other performance evaluations have been done to measure BEAGLE performance by the team behind this API [5]. Their results showcase substantial performance gains when using GPUs for partitions (sections of a genome treated as semi-independent evolutionary units). Notably, their results utilised a considerable number of site patterns. However, for lower counts of site patterns, their results suggest that using only CPU multi-threading could yield faster results. They also indicate that using multiple GPUs is not optimal unless analysing a large number of site patterns.

These observations are particularly relevant to viral genomes, whose number of site patterns is too small to exploit GPU computing, especially when partitioned. For instance, the Dengue Virus (DENV) has a genome of 11 kb, which encodes 10 genes, three structural and seven non-structural, plus a “2K fragment” between genes NS4A and NS4B [7]. When partitioned according to the proteins, the number of site patterns per partition ranges from 40 to 1,300.

An incorrect allocation of resources could slow the execution time of a BEAST X run. Furthermore, scientific computing has a substantial carbon footprint [8], so avoiding unnecessary resource use is a crucial step towards environmentally responsible research.

This manuscript aims to benchmark BEAST X, using real genomes of DENV and simulated sequences, with different BEAGLE configurations to establish guidelines for optimising running times for phylodynamic applications, particularly for genome surveillance [9] and pandemic preparedness [10]. While BEAST X benchmarks have already been done, notably by CIPRES [6], they intended to find ways of optimising cost-effectiveness, for instance, only switching to GPU computing if there is a very significant gain over CPUs. Here, we focus primarily on running time. We will only consider nucleotide substitution models, and we note that the speedup from parallelisation is more substantial for codon or amino-acid models [5].

## II. Methods

### A. Real Data

We obtained the benchmarking data with its associated metadata from the NextStrain DENV project [11] dataset, which is a curated selection of publicly available sequences from GenBank. Out of the 52,665 sequences available, we applied several filters to ensure the data met the following criteria:

- Sequences had to be extracted from human hosts.
- Sequences had to be complete genomes.
- Sequences had to have GenBank annotations specifying the location of all 11 genetic regions of interest (10 genes plus the 2K segment) to facilitate partitioning later in our pipeline.
- Given the relatively high rate of nucleotide substitution in dengue [12], sequences had to have a specified sampling month to avoid biases that could arise from uncertainty in sampling times [2].
- If the exact date of a sequence was not provided, the date had to be rounded to the 15^*th*^ of the month.

Only sequences meeting all these criteria were included in the final dataset. Table I contains serotype representation across the final dataset.

**Table I.** Serotype representation in real benchmarking data. The table shows the counts and percentages of each serotype relative to the total number of samples (376).

|  | DENV1 | DENV2 | DENV3 | DENV4 |
| --- | --- | --- | --- | --- |
| **Count** | 135 | 111 | 96 | 34 |
| **Percentage** | 35.90% | 29.52% | 25.53% | 9.04% |

From the 376 filtered sequences, we randomly sub-sampled 30 samples from each serotype to form two benchmarking datasets. We retained only the protein-coding region, and each dataset included both partitioned and non-partitioned versions. We split the partitioned sequences into 11 sections (10 genes plus the 2K protein) according to their respective GFF3 annotation files. We used Mafft (v7.490) [13] for the alignment. The final accessions are available at https://github.com/Fossam40/BeagleBeastBenchmark.

### B. Simulated sequences

We used simulated data generated with Seq-Gen [14] to control the number of site patterns and to measure how this factor impacts running times across different BEAGLE configurations. We sampled the trees from the posterior of the BEAST X runs using real data. To obtain branch lengths in units of substitutions per site (subs/site), we multiplied the initial branch lengths by a molecular clock rate. We used the *HKY* + Γ_4_ [15] model, with parameters sampled from the prior, to generate the sequences. The scripts used to generate those sequences are available in the GitHub repository https://github.com/Fossam40/BeagleBeastBenchmark with the XML files associated with the benchmarks.

### C. Bayesian inference parameters

All the benchmarks were done under BEAST X v1.10.5 (beta 5, commit 1d511b1), using *HKY* + Γ_4_ [15] as the substitution model, a constant-size coalescent tree prior, and a relaxed molecular clock model with an underlying lognormal distribution [16]. This molecular clock model was used since DENV has been found to follow a molecular clock with some rate differences across genotypes [12]. We used the default priors suggested by the graphical interface BEAUTi [17]. Additionally, we set the chain length to 10^8^ and recorded parameters every 10^4^ steps.

We left both the clock and substitution models unlinked to account for possible differences in fitness pressure across genes. The XML files for all runs are available at https://github.com/Fossam40/BeagleBeastBenchmark.

The benchmark runs predate the full release of BEAST X v10.5.0, so the fifth beta of the version was used, with the last release of BEAGLE (v4.0.1). The Java version was 22.3.1.

### D. Benchmarking methods

After defining the same model for all our runs (available as an XML file in the GitHub repository), we evaluated different configurations of the following flags: `-beagle_GPU`, `-threads`, `-beagle_instances`, and `-beagle_SSE` for both partitioned and non-partitioned real data. Our focus was on the impact of different CPU/GPU configurations. The various configurations of the running parameters are detailed in Supplementary Table IV. A comprehensive list of all parameters used is available on the associated GitHub page.

To compare CPU and GPU performance on simulated sequences, we ran tests comprising four consecutive runs, using one core, two cores, four cores, and one GPU. With this method, we obtain running times under roughly the same conditions, which are then normalised by comparing them to the single-core run. A similar method was applied to compare single-and double-GPU runs, this time normalising by the running time of the single-GPU test.

All runs were conducted in the HPC Core Facility of the Institut Pasteur in Paris, France. Each run was replicated at least two times to account for variability due to other jobs running on the same node. The processors of each computing node were identical (AMD EPYC 7552 48-Core Processor), and for GPU runs, we utilised Nvidia A40 cards.

## III. Results

Table II presents the running times in hours for the two datasets, with different configurations of partitioning, threads, subpartitions, and GPUs. Each row corresponds to a different experiment, with its configurations and results. The fastest way to run BEAST with these datasets is shown in experiments three and nine, while the slowest were experiments two and seven, for partitioned and non-partitioned versions, respectively. While most experiments have standard deviations under 2.5, experiment seven has a 5-hour standard deviation for both datasets, and 6 hours for dataset 2. Each real-data experiment was replicated at least three times, which is why the average running times are reported.

**Table II.**
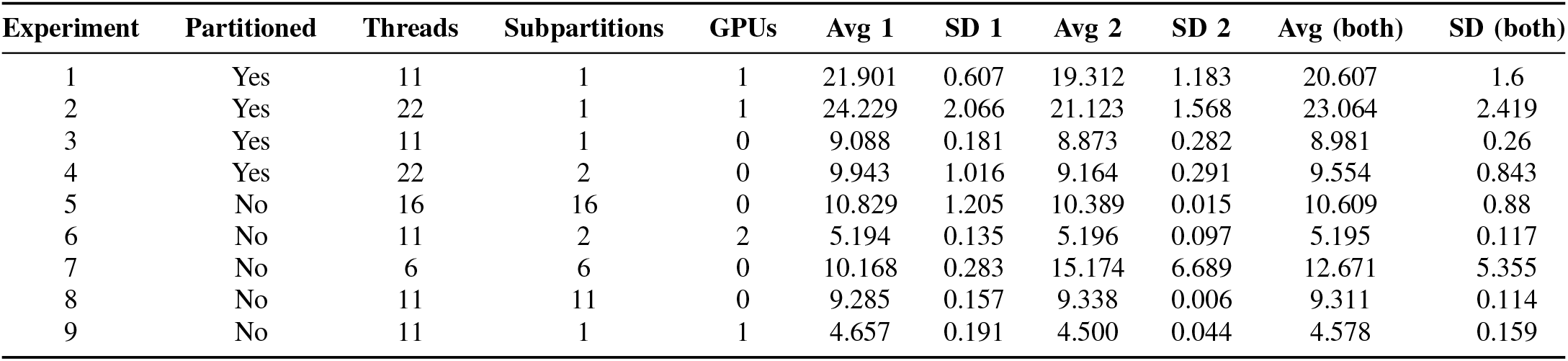
Running times in hours of datasets 1 and 2. Dataset 1 has an average of 480 site patterns per partition, while dataset 2 has an average of 493 site patterns per partition.

| Experiment | Partitioned | Threads | Subpartitions | GPUs | Avg 1 | SD 1 | Avg 2 | SD 2 | Avg (both) | SD (both) |
| --- | --- | --- | --- | --- | --- | --- | --- | --- | --- | --- |
| 1 | Yes | 11 | 1 | 1 | 21.901 | 0.607 | 19.312 | 1.183 | 20.607 | 1.6 |
| 2 | Yes | 22 | 1 | 1 | 24.229 | 2.066 | 21.123 | 1.568 | 23.064 | 2.419 |
| 3 | Yes | 11 | 1 | 0 | 9.088 | 0.181 | 8.873 | 0.282 | 8.981 | 0.26 |
| 4 | Yes | 22 | 2 | 0 | 9.943 | 1.016 | 9.164 | 0.291 | 9.554 | 0.843 |
| 5 | No | 16 | 16 | 0 | 10.829 | 1.205 | 10.389 | 0.015 | 10.609 | 0.88 |
| 6 | No | 11 | 2 | 2 | 5.194 | 0.135 | 5.196 | 0.097 | 5.195 | 0.117 |
| 7 | No | 6 | 6 | 0 | 10.168 | 0.283 | 15.174 | 6.689 | 12.671 | 5.355 |
| 8 | No | 11 | 11 | 0 | 9.285 | 0.157 | 9.338 | 0.006 | 9.311 | 0.114 |
| 9 | No | 11 | 1 | 1 | 4.657 | 0.191 | 4.500 | 0.044 | 4.578 | 0.159 |

The “Experiment” header is a numbering of the rows. The “Partitioned” header indicates whether the dataset was partitioned by genes. The “Threads” header shows the number of threads that were made available to BEAST X, which we controlled with the “-threads” option, while “GPUs” is the number of allocated GPUs (zero for CPU-only runs). The subpartition header indicates the number of BEAGLE instances per partition. The individual runs, with additional details, are listed in Table V. The number of site patterns per partition is available in Table IV.

Figure 1 is a bar plot of the results of Table II. The best option for the complete dataset is to use a single GPU, which yields an almost two-fold increase in performance compared to CPU runs. However, using two GPUs is slower than using one. Increasing the number of CPU threads from 6 to 11 improved performance, but further increasing to 16 degraded it.

**Figure 1.**
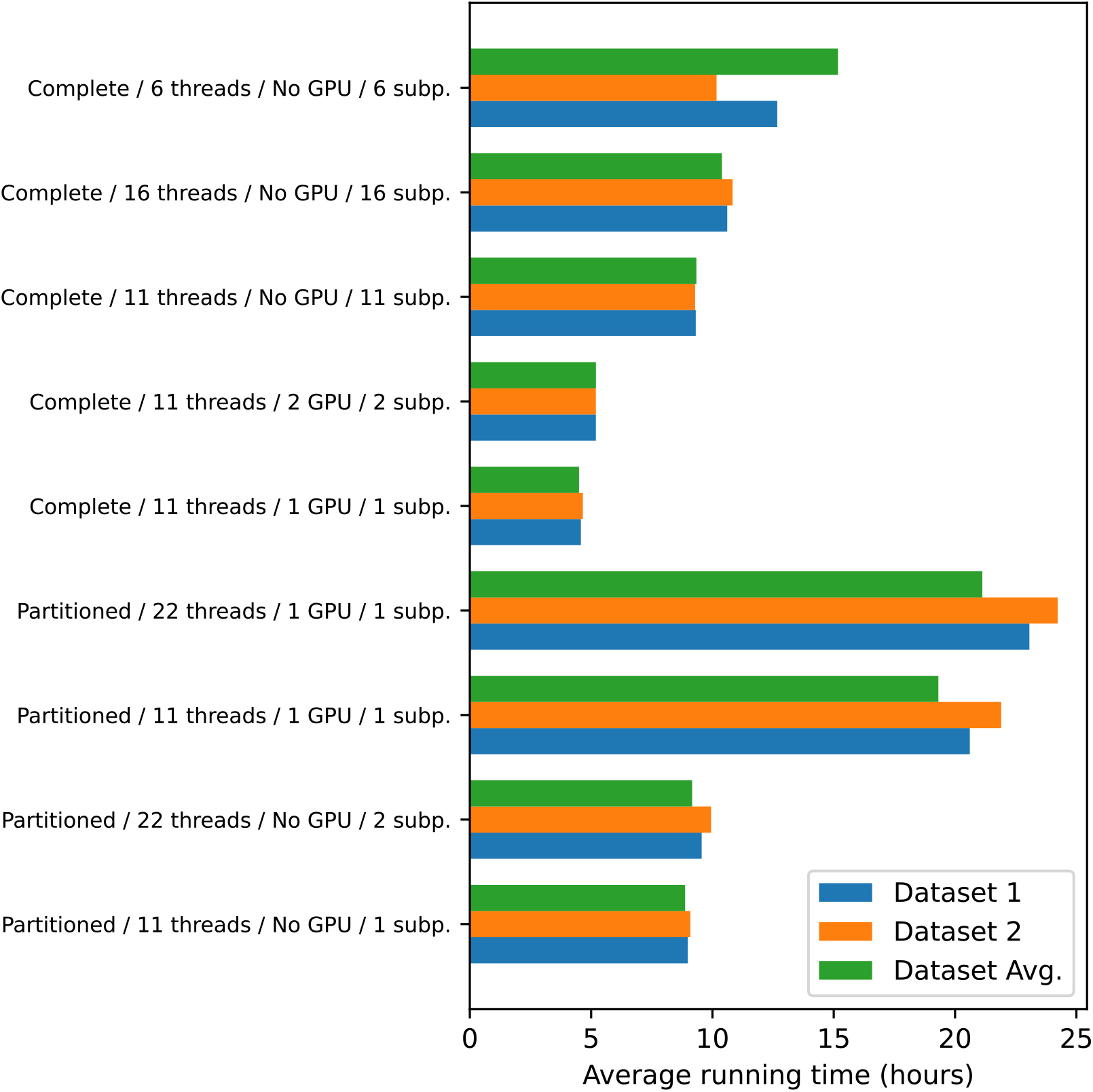
Average running times in hours for all datasets and parameters. Values are sorted in decreasing order, separating partitioned and non-partitioned.

For the partitioned versions of the datasets, GPU runs are more than twice as slow as multi-threading approaches. Using a thread per partition was the fastest option, performing on average slightly better than assigning two threads per partition.

Figure 2 shows the relative speedup of using two cores, four cores and a GPU against the number of site patterns, as measured in the artificial data. While GPU runs were slower than single-core runs for 665 site patterns, they became faster than multi-core runs for more than 860 site patterns. A detailed view of the individual runs is shown in Table VI.

**Figure 2.**
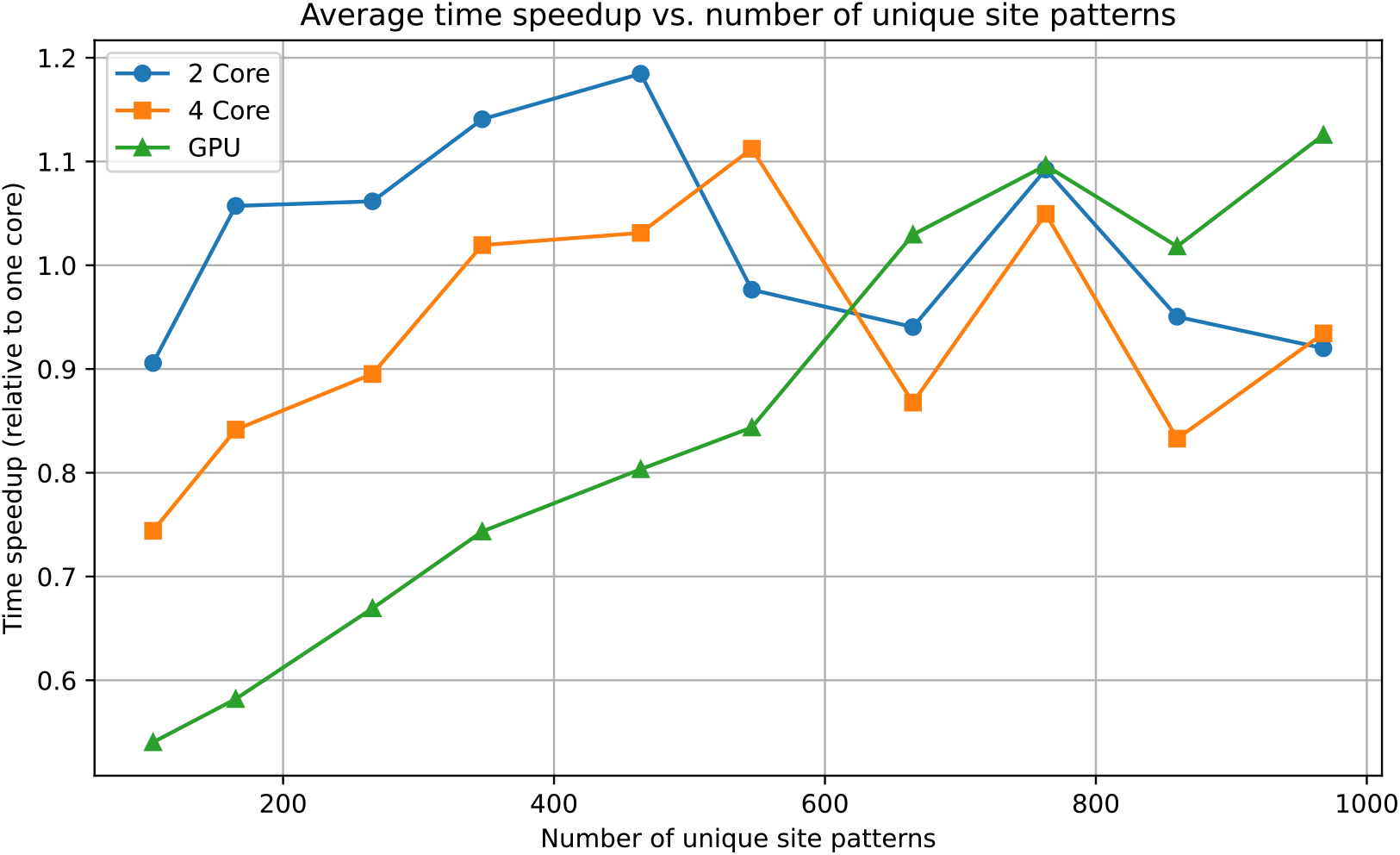
Average speedup factor for multi-core (two and four cores) and GPU runs compared to single-core runs, depending on the number of site patterns. Measurements were done in increments of around 100 site patterns.

Similarly, Figure 3 shows the average relative speedup of using two GPUs over using one. Single-GPU runs were, on average, faster up to around 25000 site patterns, after which using two GPUs yielded minor improvements. The details of every run can be found in Table VII.

**Figure 3.**
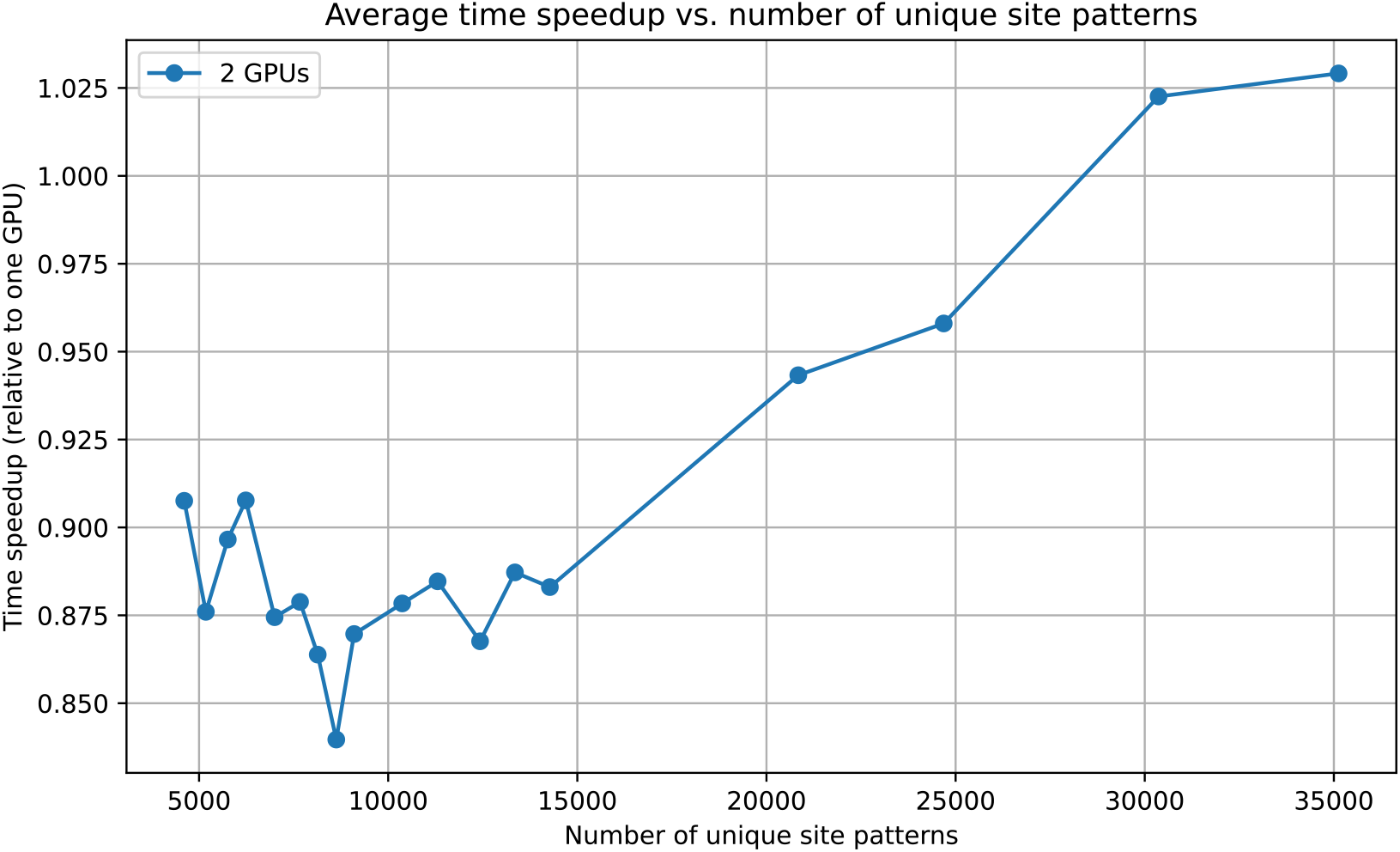
Average speedup factor using two NVIDIA A40 GPUs compared to using one NVIDIA A40. Measurements were performed in increments of approximately 500 site patterns up to 14274, and then in increments of 5000 to show higher counts.

## Discussion

Our tests with real DENV data indicate that, for the complete datasets 1 and 2, using a single GPU yields an almost two-fold performance increase compared to CPU runs. For partitioned datasets, GPU runs are more than twice as slow as multi-threading approaches, with assigning one thread per partition being the fastest option, slightly outperforming two threads per partition.

The simulated data runs suggest that, for an Nvidia A40 card, switching from CPU to GPU is recommended only for more than 860 sites. Under that threshold, one should consider using only CPU cores with some level of multithreading. Although benchmarks with two GPUs were faster than single-GPU runs with 30,000 site patterns, the speedup was not significant enough to justify the higher economic and environmental costs of using an extra GPU. More tests with higher site patterns are needed to determine when multi-GPU approaches become truly competitive.

Our findings align with the CIPRES benchmark [6], which also showed diminishing returns with excessive threading. However, our results with BEAST X v1.10.5 and modern hardware (NVIDIA A40 GPUs, AMD EPYC processors) provide updated guidelines for current computational infrastructures. Unlike the CIPRES results, which focused on cost-effectiveness and were conducted with BEAST version 1.6.1 over a decade ago, our GPU threshold of 860 site patterns provides a clear performance-based guideline optimised for contemporary hardware configurations.

Some benchmarking runs exhibited anomalous run-times, despite the output files being normal. That is, the posterior distribution was sampled correctly according to the MCMC trace, despite differences in runtimes. This is the case for benchmark number 50 in Table V, which ran for more than twice as long as other runs with the same parameter. This results in a substantial standard deviation in experiment seven of Table II. Experiment two in the same table also showed substantial variation across runs. We attempted to compensate for this by increasing the number of runs, without success. Our main hypothesis is that benchmark number 50 was affected by high job demand in the cluster at the time, resulting in decreased overall performance, but the source of the variation observed in experiment two in Table II remains unclear.

Our benchmarks do not measure how different BEAST models affect the performance. We hypothesise that the number of site patterns required to justify parameter changes varies across models. One possible method to predict the performance of a BEAST run, regardless of the models used, could be to run the MCMC chain for a short period, and use the time per million states to evaluate performance. Further research would be needed to determine how long this sample run should last for the test to be meaningful.

Our results indicate that hardware allocation can drastically impact running times. Furthermore, there is no universal configuration that is optimal for any dataset, so one must account for dataset characteristics (e.g., the number of site patterns across partitions) when choosing BEAGLE settings.

## Acronyms

API: Application Programming Interface.
BEAGLE: Broad-platform Evolutionary Analysis General Likelihood Evaluator.
BEAST: Bayesian Evolutionary Analysis Sampling.
BEAUTi: Bayesian Evolutionary Analysis Utility.
DENV: Dengue Virus.
GPU: Graphics Processing Unit.
HPC: High-Performance Computing.
MCMC: Markov Chain Monte Carlo.

## IV. Supplements

**Table IV.**
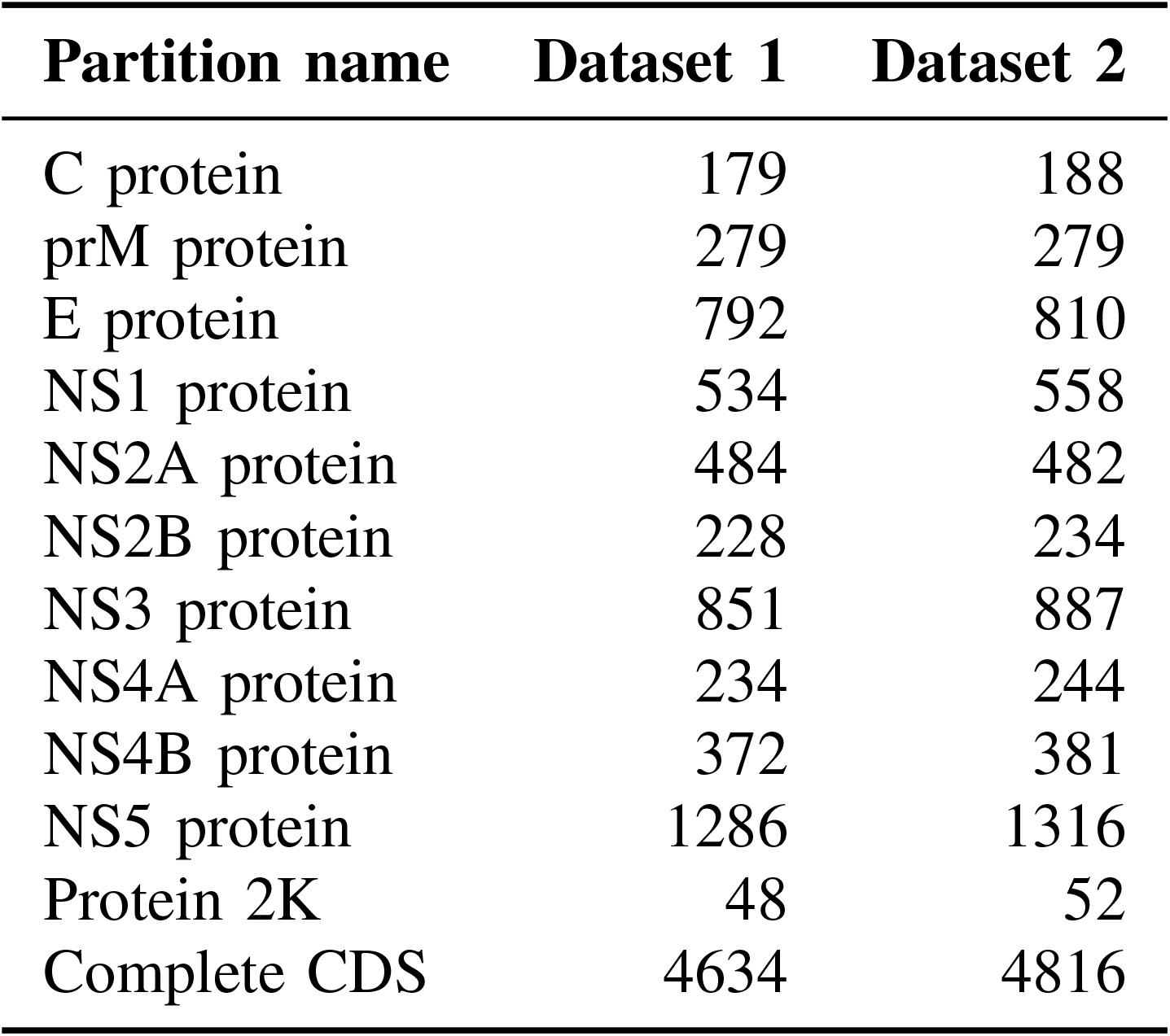
Site patterns per partition. Every partition (except the last one) corresponds to a protein.

| Partition name | Dataset 1 | Dataset 2 |
| --- | --- | --- |
| C protein | 179 | 188 |
| prM protein | 279 | 279 |
| E protein | 792 | 810 |
| NS1 protein | 534 | 558 |
| NS2A protein | 484 | 482 |
| NS2B protein | 228 | 234 |
| NS3 protein | 851 | 887 |
| NS4A protein | 234 | 244 |
| NS4B protein | 372 | 381 |
| NS5 protein | 1286 | 1316 |
| Protein 2K | 48 | 52 |
| Complete CDS | 4634 | 4816 |

**Table V.** Information on all runs, with memory allocated (in GB) in the cluster and their BEAST X options. The `-beagle_SSE` flag was always accompanied by the `-beagle_CPU` flag. The number of allocated cores matches the value specified by the-threads option. Some runs were allocated less memory than others, but memory usage was below 25% in all cases. The CPU used was an AMD EPYC 7552 48-Core Processor.

| Row | Dataset | Partitioned | -beagle_GPU | -threads | -beagle_instances | -beagle_SSE | Mem | Time in (days-)h:m:s |
| --- | --- | --- | --- | --- | --- | --- | --- | --- |
| 1 | 1 | Yes | Yes | 11 | No | No | 20 | 21:06:57 |
| 2 | 1 | Yes | Yes | 11 | No | No | 20 | 21:59:36 |
| 3 | 1 | Yes | Yes | 11 | No | No | 20 | 22:35:43 |
| 4 | 1 | Yes | Yes | 22 | No | No | 20 | 1-01:27:52 |
| 5 | 1 | Yes | Yes | 22 | No | No | 20 | 1-01:31:54 |
| 6 | 1 | Yes | Yes | 22 | No | No | 20 | 20:39:45 |
| 7 | 1 | Yes | Yes | 22 | No | No | 20 | 23:10:59 |
| 8 | 1 | Yes | Yes | 22 | No | No | 20 | 1-02:18:19 |
| 9 | 1 | Yes | No | 11 | No | Yes | 20 | 09:07:32 |
| 10 | 1 | Yes | No | 11 | No | Yes | 20 | 09:17:20 |
| 11 | 1 | Yes | No | 11 | No | Yes | 20 | 08:51:02 |
| 12 | 1 | Yes | No | 22 | 2 | Yes | 20 | 09:06:40 |
| 13 | 1 | Yes | No | 22 | 2 | Yes | 20 | 11:22:27 |
| 14 | 1 | Yes | No | 22 | 2 | Yes | 20 | 09:20:42 |
| 15 | 1 | No | No | 16 | 16 | Yes | 20 | 09:54:58 |
| 16 | 1 | No | No | 16 | 16 | Yes | 20 | 10:02:18 |
| 17 | 1 | No | No | 16 | 16 | Yes | 20 | 12:31:54 |
| 18 | 1 | No | No<sup>a</sup> | 11 | 2 | No | 20 | 05:00:43 |
| 19 | 1 | No | No<sup>a</sup> | 11 | 2 | No | 20 | 05:14:07 |
| 20 | 1 | No | No<sup>a</sup> | 11 | 2 | No | 20 | 05:20:04 |
| 21 | 1 | No | No | 6 | 6 | Yes | 20 | 10:19:51 |
| 22 | 1 | No | No | 6 | 6 | Yes | 20 | 10:24:13 |
| 23 | 1 | No | No | 6 | 6 | Yes | 20 | 09:46:13 |
| 24 | 1 | No | No | 11 | 11 | Yes | 20 | 09:06:20 |
| 25 | 1 | No | No | 11 | 11 | Yes | 20 | 09:29:15 |
| 26 | 1 | No | No | 11 | 11 | Yes | 20 | 09:15:45 |
| 27 | 1 | No | Yes | 11 | No | No | 20 | 04:52:45 |
| 28 | 1 | No | Yes | 11 | No | No | 20 | 04:24:47 |
| 29 | 1 | No | Yes | 11 | No | No | 20 | 04:40:42 |
| 30 | 2 | Yes | Yes | 11 | No | No | 5 | 18:27:32 |
| 31 | 2 | Yes | Yes | 11 | No | No | 5 | 20:59:07 |
| 32 | 2 | Yes | Yes | 11 | No | No | 5 | 18:29:30 |
| 33 | 2 | Yes | Yes | 22 | No | No | 20 | 23:01:22 |
| 34 | 2 | Yes | Yes | 22 | No | No | 20 | 21:09:47 |
| 35 | 2 | Yes | Yes | 22 | No | No | 20 | 19:10:57 |
| 36 | 2 | Yes | No | 11 | No | Yes | 5 | 09:05:28 |
| 37 | 2 | Yes | No | 11 | No | Yes | 5 | 08:28:27 |
| 38 | 2 | Yes | No | 11 | No | Yes | 5 | 09:03:12 |
| 39 | 2 | Yes | No | 22 | 2 | Yes | 5 | 08:45:10 |
| 40 | 2 | Yes | No | 22 | 2 | Yes | 5 | 09:22:11 |
| 41 | 2 | Yes | No | 22 | 2 | Yes | 5 | 09:22:11 |
| 42 | 2 | No | No | 16 | 16 | Yes | 5 | 10:24:24 |
| 43 | 2 | No | No | 16 | 16 | Yes | 5 | 10:22:16 |
| 44 | 2 | No | No | 16 | 16 | Yes | 5 | 10:23:17 |
| 45 | 2 | No | No<sup>a</sup> | 11 | 2 | No | 5 | 05:06:14 |
| 46 | 2 | No | No<sup>a</sup> | 11 | 2 | No | 5 | 05:19:47 |
| 47 | 2 | No | No<sup>a</sup> | 11 | 2 | No | 5 | 05:09:19 |
| 48 | 2 | No | No | 6 | 6 | Yes | 5 | 10:17:48 |
| 49 | 2 | No | No | 6 | 6 | Yes | 5 | 10:35:41 |
| 50 | 2 | No | No | 6 | 6 | Yes | 5 | 1-00:37:55 |
| 51 | 2 | No | No | 11 | 11 | Yes | 5 | 09:20:45 |
| 52 | 2 | No | No | 11 | 11 | Yes | 5 | 09:19:54 |
| 53 | 2 | No | No | 11 | 11 | Yes | 20 | 09:20:07 |
| 54 | 2 | No | Yes | 11 | No | No | 20 | 04:33:21 |
| 55 | 2 | No | Yes | 11 | No | No | 20 | 04:26:59 |
| 56 | 2 | No | Yes | 11 | No | No | 20 | 04:29:35 |

<sup>a</sup>Run used `-beagle_order 1,2` to allocate GPUs

**Table VI.**
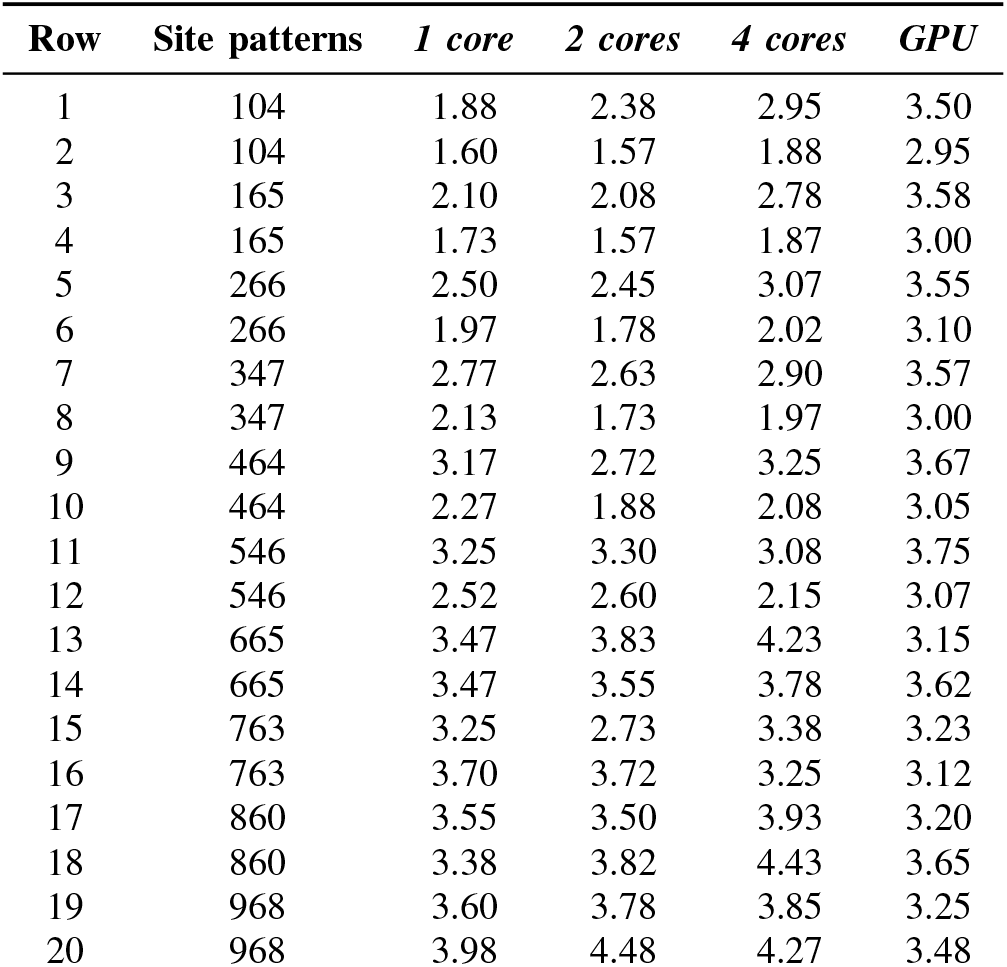
Information on all simulated sequence runs comparing CPU and GPU. The number of site patterns is specified, as are the running times (in hours) of subsequent benchmarks on one core, two cores, four cores, and the GPU. All runs were allocated 5GB of memory. The CPU used was an AMD EPYC 7552 48-Core Processor.

| Row | Site patterns | 1 core | 2 cores | 4 cores | GPU |
| --- | --- | --- | --- | --- | --- |
| 1 | 104 | 1.88 | 2.38 | 2.95 | 3.50 |
| 2 | 104 | 1.60 | 1.57 | 1.88 | 2.95 |
| 3 | 165 | 2.10 | 2.08 | 2.78 | 3.58 |
| 4 | 165 | 1.73 | 1.57 | 1.87 | 3.00 |
| 5 | 266 | 2.50 | 2.45 | 3.07 | 3.55 |
| 6 | 266 | 1.97 | 1.78 | 2.02 | 3.10 |
| 7 | 347 | 2.77 | 2.63 | 2.90 | 3.57 |
| 8 | 347 | 2.13 | 1.73 | 1.97 | 3.00 |
| 9 | 464 | 3.17 | 2.72 | 3.25 | 3.67 |
| 10 | 464 | 2.27 | 1.88 | 2.08 | 3.05 |
| 11 | 546 | 3.25 | 3.30 | 3.08 | 3.75 |
| 12 | 546 | 2.52 | 2.60 | 2.15 | 3.07 |
| 13 | 665 | 3.47 | 3.83 | 4.23 | 3.15 |
| 14 | 665 | 3.47 | 3.55 | 3.78 | 3.62 |
| 15 | 763 | 3.25 | 2.73 | 3.38 | 3.23 |
| 16 | 763 | 3.70 | 3.72 | 3.25 | 3.12 |
| 17 | 860 | 3.55 | 3.50 | 3.93 | 3.20 |
| 18 | 860 | 3.38 | 3.82 | 4.43 | 3.65 |
| 19 | 968 | 3.60 | 3.78 | 3.85 | 3.25 |
| 20 | 968 | 3.98 | 4.48 | 4.27 | 3.48 |

**Table VII.**
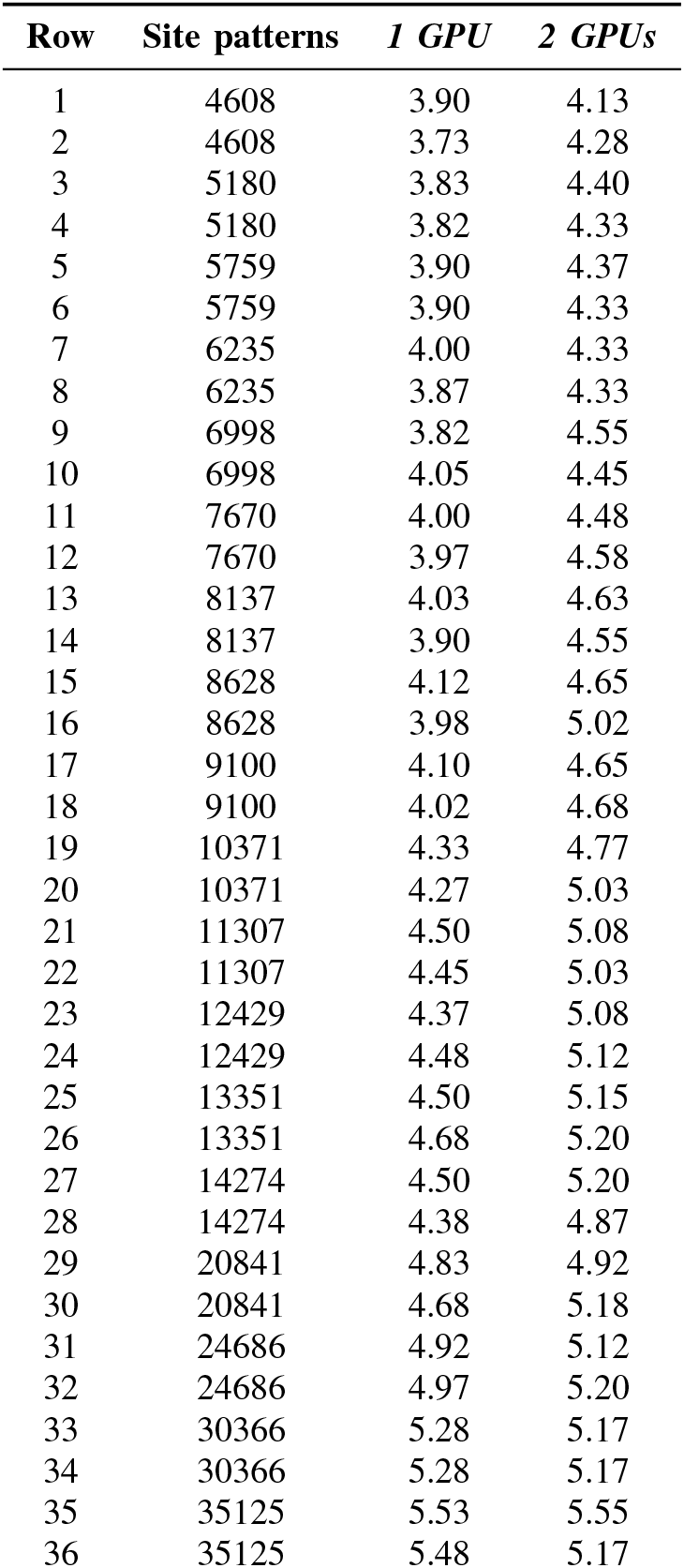
Information on all simulated sequence runs comparing a single GPU and two GPUs. The number of site patterns is specified, as are the running times (in hours) of subsequent benchmarks on one core, two cores, four cores, and the GPU. All runs were allocated 5GB of memory.

| Row | Site patterns | 1 GPU | 2 GPUs |
| --- | --- | --- | --- |
| 1 | 4608 | 3.90 | 4.13 |
| 2 | 4608 | 3.73 | 4.28 |
| 3 | 5180 | 3.83 | 4.40 |
| 4 | 5180 | 3.82 | 4.33 |
| 5 | 5759 | 3.90 | 4.37 |
| 6 | 5759 | 3.90 | 4.33 |
| 7 | 6235 | 4.00 | 4.33 |
| 8 | 6235 | 3.87 | 4.33 |
| 9 | 6998 | 3.82 | 4.55 |
| 10 | 6998 | 4.05 | 4.45 |
| 11 | 7670 | 4.00 | 4.48 |
| 12 | 7670 | 3.97 | 4.58 |
| 13 | 8137 | 4.03 | 4.63 |
| 14 | 8137 | 3.90 | 4.55 |
| 15 | 8628 | 4.12 | 4.65 |
| 16 | 8628 | 3.98 | 5.02 |
| 17 | 9100 | 4.10 | 4.65 |
| 18 | 9100 | 4.02 | 4.68 |
| 19 | 10371 | 4.33 | 4.77 |
| 20 | 10371 | 4.27 | 5.03 |
| 21 | 11307 | 4.50 | 5.08 |
| 22 | 11307 | 4.45 | 5.03 |
| 23 | 12429 | 4.37 | 5.08 |
| 24 | 12429 | 4.48 | 5.12 |
| 25 | 13351 | 4.50 | 5.15 |
| 26 | 13351 | 4.68 | 5.20 |
| 27 | 14274 | 4.50 | 5.20 |
| 28 | 14274 | 4.38 | 4.87 |
| 29 | 20841 | 4.83 | 4.92 |
| 30 | 20841 | 4.68 | 5.18 |
| 31 | 24686 | 4.92 | 5.12 |
| 32 | 24686 | 4.97 | 5.20 |
| 33 | 30366 | 5.28 | 5.17 |
| 34 | 30366 | 5.28 | 5.17 |
| 35 | 35125 | 5.53 | 5.55 |
| 36 | 35125 | 5.48 | 5.17 |

